# VPS8D, a CORVET subunit, is required to maintain the contractile vacuole complex in *Tetrahymena thermophila*

**DOI:** 10.1101/2023.11.07.566071

**Authors:** Chao-Yin Cheng, Josefina Hernández, Aaron P. Turkewitz

**Affiliations:** Department of Molecular Genetics and Cell Biology, University of Chicago, Chicago, IL, USA

**Keywords:** *Tetrahymena*, contractile vacuole complex, CORVET (class C **cor**e vacuole/**e**ndosome-**t**ethering), RNA hairpin, osmotic sensitivity, Ciliate

## Abstract

Contractile vacuole complexes (CVCs) are complex osmoregulatory organelles, with vesicular (bladder) and tubular (spongiome) subcompartments. The mechanisms that underlie their formation and maintenance within the eukaryotic endomembrane network are poorly understood. In the Ciliate *Tetrahymena thermophila*, six differentiated CORVETs (class C core vacuole/endosome tethering complexes), with Vps8 subunits designated A-F, are likely to direct endosomal trafficking. Vps8Dp localizes to both bladder and spongiome. We show by inducible knockdown that *VPS8D* is essential to CVC organization and function. *VPS8D* knockdown increased susceptibility to osmotic shock, tolerated in the wildtype but triggering irreversible lethal swelling in the mutant. The knockdown rapidly triggered contraction of the spongiome and lengthened the period of the bladder contractile cycle. More prolonged knockdown resulted in disassembly of both the spongiome and bladder, and dispersal of proteins associated with those compartments. In stressed cells where the normally singular bladder is replaced by numerous vesicles bearing bladder markers, Vps8Dp concentrated conspicuously at long-lived inter-vesicle contact sites, consistent with tethering activity. Similarly, Vps8Dp in cell-free preparations accumulated at junctions formed after vacuoles came into close contact. Also consistent with roles for Vps8Dp in tethering and/or fusion were the emergence in knockdown cells of multiple vacuole-related structures, replacing the single bladder.

**Synopsis:** In the Ciliate Tetrahymena thermophila, *VPS8D*, which encodes a subunit of a non-conventional CORVET complex, is an essential determinant of the contractile vacuole complex (CVC). VPS8D knockdown results in retraction and dispersal of the spongiome, and disappearance of the bladder, reinforcing the view that CVCs arise from endosomal trafficking. Intermediate knockdown phenotypes and Vps8Dp localization support a role in homotypic tethering.

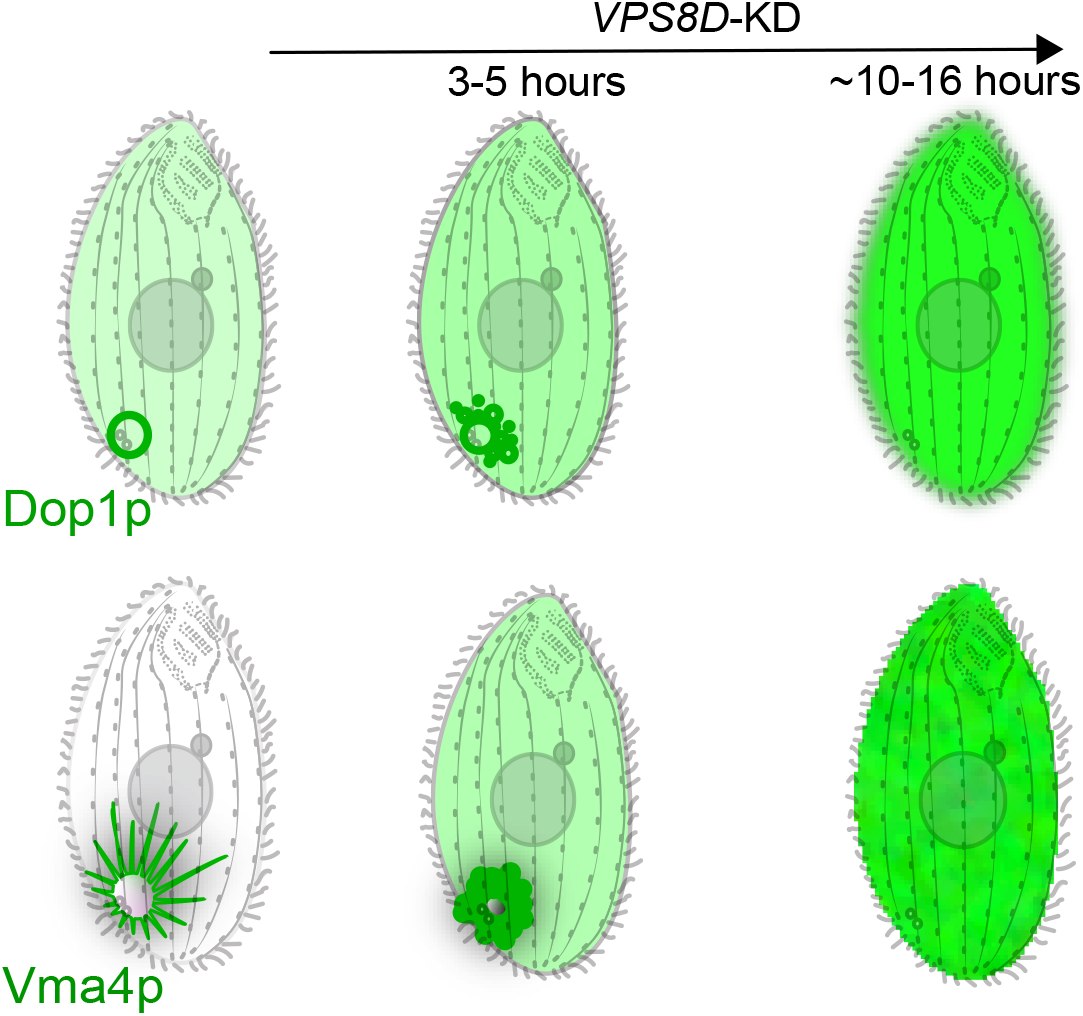

## Introduction

Contractile vacuole complexes (CVCs) are a morphologically diverse set of organelles that are found very broadly throughout eukaryotes, and particularly in eukaryotic protists. Their recognized role is in osmoregulation, a critical issue for single-celled organisms inhabiting freshwater and therefore hyperosmotic environments because constant passive water uptake across the plasma membrane threatens cellular via viability, including maintaining physiological cytoplasmic ion concentrations (Naitoh et al., 1997, Tominaga et al., 1998a, Klauke and Plattner, 2000, Kissmehl et al., 2002, Tani et al., 2002, Reuter et al., 2013). Notwithstanding their heterogeneity, CVCs all appear built on similar organizing principles. They are comprised of one or more large vacuolar compartments that function as water-storing bladders, that are ringed by a network of tubular extensions called the spongiome (Allen and Naitoh, 2002). The existing data are consistent with the general idea that cytoplasmic water crosses the spongiome membranes via aquaporins or similar channels and driven by gradients dependent in part on proton-pumping V^+^-ATPases. Water in the spongiome lumen diffuses to swell the connected bladder. At some point, the bladder membrane fuses with but is not integrated into the plasma membrane, and the soluble contents are rapidly propelled outside the cell by dramatic contraction of the bladder. This membrane fusion event in the Amoebozoan *Dictyostelium discoideum* is regulated at least in part by triggered calcium release from the bladder lumen, which acts to downregulate a CVC-specific Rab (Parkinson et al., 2014). More generally, ion mobilization from the CVC lumen and/or excretion during bladder contraction may play homeostatic and/or signalling roles (Ladenburger et al., 2006, Malchow et al., 2008, Ladenburger et al., 2009, Patel and Docampo, 2010, Sivaramakrishnan and Fountain, 2012). Following bladder contraction, the fusion pore to the plasma membrane reseals and the bladder is ready for re-filling. Though functionally similar, the precise structures involved in this cycle and their dynamic interactions may differ in important details between lineages. For example, in the Ciliate Paramecium the reticular spongiome is comprised of two distinct zones, and the contractile cycle includes membrane fission and fusion between the spongiome and bladder (Allen, 2000), neither of which features in the CVC of the Kinetoplastid *Trypanosoma cruzi* (Jimenez et al., 2022).

CVCs are not found in yeasts or other lineages in which cells secrete and assemble external walls that can counterbalance osmotic pressures with physical restraints, nor in multicellular organisms like animals where the tonicity of the extracellular milieu is under organismal control. Since the most commonly employed model organisms belong to these groups, CVCs have been relatively understudied and consequently many central issues remain unsettled. One largely open question is how the CVC is integrated with other pathways of eukaryotic membrane trafficking, including the mechanisms that underlie its maintenance in vegetative cells and inheritance during cell division. A number of proteins whose homologs are involved in membrane trafficking in fungi and animals, have been physically and/or functionally associated with the CVC in diverse eukaryotic lineages. Early work in *D. discoideum* linked both and the clathrin assembly factor AP180 to CVC function, suggesting a putative connection between endocytic trafficking and CVC maintenance (Wen et al., 2009, Macro et al., 2012). The CVC is nonetheless normally isolated from bulk endocytic trafficking, in both *Dictyostelium* and the Ciliate *Tetrahymena thermophila*, based on studies using endocytic tracers (Gabriel et al., 1999, Cheng et al., 2023). Whether the same trafficking pathways maintain the CVC in all lineages remains an open question. A related but more far-reaching question is whether CVCs emerged independently in different lineages, or whether they stem from a single ancestral origin. The fact that all known CVCs rely on vacuolar ATPases might argue for the latter, but vacuolar ATPases function in a very broad range of compartments in the endomembrane network (Stevens and Forgac, 1997).

In addition to clathrin and AP180, other endosomal protein homologs were also shown to be physically and/or functionally associated with the CVC in Dictyostelium, including AP2, epsin, the endosomal synaptobrevin homolog Vamp7B, Rabs 11 and 8, and an amphiphysin (I-BAR) family protein (Harris et al., 2001, Stavrou and O’Halloran, 2006, Wen et al., 2009, Linkner et al., 2014, Manna et al., 2023, Du et al., 2008, Essid et al., 2012, Macro et al., 2012). Homologs for a subset of these, including a Rab11 family member, were subsequently found associated with the CVC in *Trypanosoma cruzi* (Ulrich et al., 2011). The Ciliates are a third, very distantly related lineage with prominent CVCs, about which relatively little is known about membrane trafficking determinants. Comprehensive localization surveys of SNAREs (in *Paramecium tetraaurelia*) and of Rabs (in *T. thermophila*) provided only ambiguous evidence addressing whether the CVC is an endosome-related compartment, since in both protein families the most prominent CVC-localized members did not belong to conserved endosomal subgroups but instead either belonged to other identifiable subgroups or were lineage-restricted (Bright et al., 2010, Plattner, 2013, Schonemann et al., 2013). A possible exception came from the observation that several *T. thermophila* Rabs belonging to conserved endocytic/endosomal subgroups showed secondary localization to the CVC, but with unknown functional significance (Bright et al., 2010). In contrast, strong evidence of an endosomal connection in ciliates came from analysis of *T. thermophila* Dop1p, a member of the DOPEY family of proteins that serve endosome-related roles in other lineages. In *T. thermophila*, Dop1p localizes strongly to the CVC where it is required for normal bladder discharge (Cheng et al., 2016, Cheng et al., 2023).

In this manuscript, we present analysis of *T. thermophila* VPS8D, a subunit of one of six *T. thermophila* CORVET (class C **cor**e vacuole/**e**ndosome-**t**ethering) complexes. In fungi, animals, and plants, CORVET and the related HOPS (**ho**motypic fusion and **p**rotein **s**orting) complexes are key determinants in endo-lysosomal trafficking (van der Beek et al., 2019). Current data support models in which the hetero-hexameric complexes tether two vesicle membranes by binding Rab GTPases in each of the adjoining membranes, and catalyze fusion by interacting with SNAREs (Baker et al., 2015). While many organisms have one CORVET and one HOPS, in the lineage leading to *T. thermophila* HOPS was lost and CORVET expanded to a family of six complexes, each with a unique subunit composition and cellular localization (Sparvoli et al., 2018). One of these complexes, which uniquely contains the subunit Vpd8D, was localized in an initial survey to the CVC, and subsequently shown to be present in a punctate distribution at both the bladder and spongiome, in dynamic equilibrium with a cytosolic pool (Sparvoli et al., 2020, Cheng et al., 2023). Attempts to study the function of Vpd8Dp by gene knockout were unsuccessful, since unlike the Vps8 subunits in other CORVET complexes the *VPS8D* paralog appeared to be essential (Sparvoli et al., 2018). Here we have therefore taken a different approach, using a gene-specific RNA hairpin to partially deplete the *VPS8D* transcript. The knockdown cells show a variety of CVC-associated phenotypes. The severity of the defects appears to correlate with the intensity of the knockdown, and at the extreme manifests as complete loss of both the bladder and spongiome. Both our functional and localization data are consistent with the idea that in *T. thermophila* a ciliate-adapted CORVET complex directs critical membrane tethering and fusion cycles in the CVC.

## Results

A cartoon illustrating the anatomy of the CVC in *T. thermophila*, together with a small set of proteins whose distributions were determined by *in vivo* localization studies of endogenously tagged genes, is shown in Fig. 1A. As previously reported (Sparvoli et al., 2020, Cheng et al., 2023), the CORVET subunit Vps8D, here endogenously tagged with mCherry, localizes to both the bladder and spongiome (Fig. 1A, right). Similarly, endogenously tagged CORVET subunit Vps11p localizes to the CVC, but also to other cellular loci as expected since it is a subunit of multiple CORVET complexes in this organism (Sparvoli et al., 2020) (Fig. S1).

**Figure 1:**
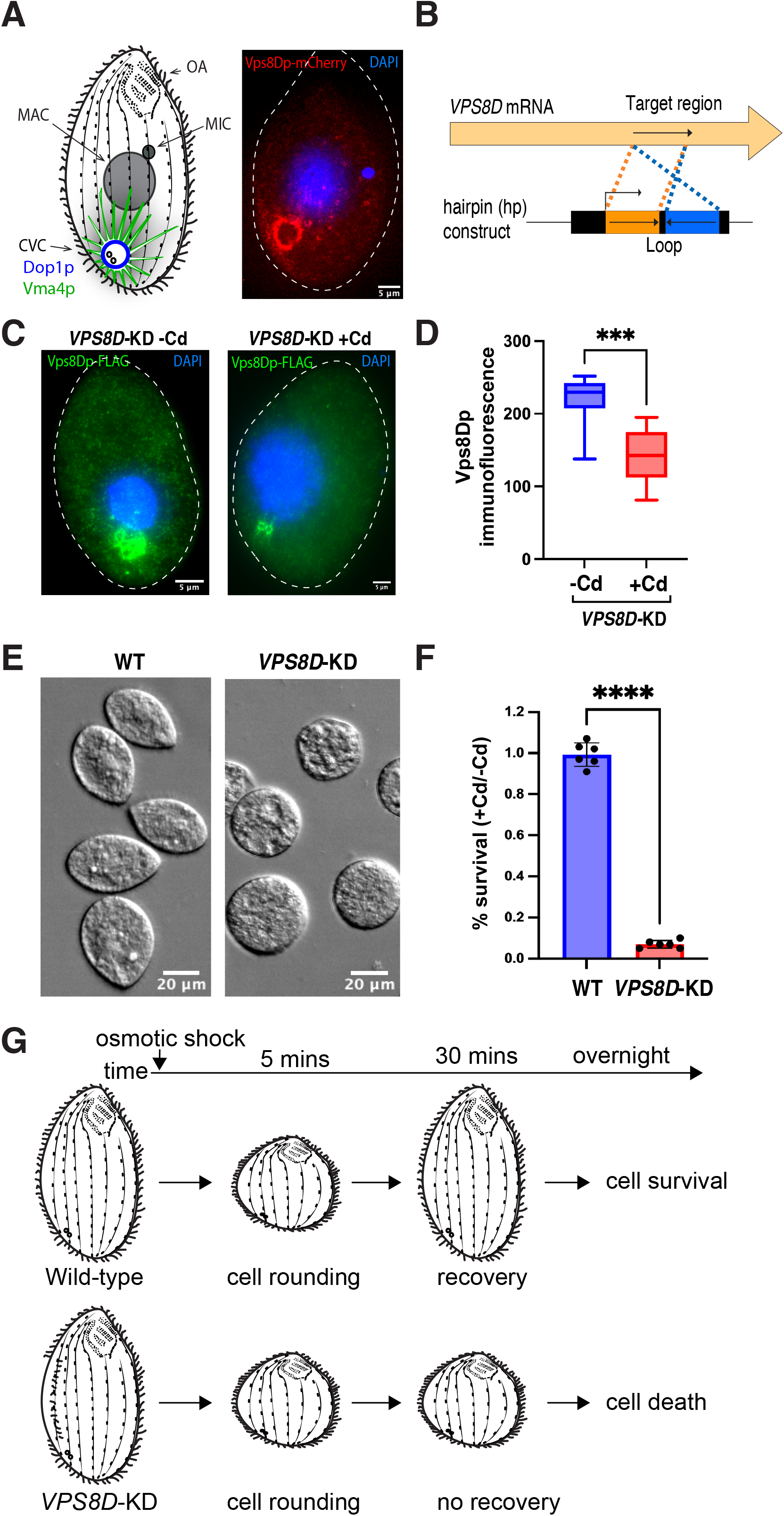
*VPS8D* plays an important role in cellular resistance to osmotic stress. A, left: CVC anatomy in *T. thermophila*. The cartoon shows previously established features of the Tetrahymena CVC including one central bladder near the cell posterior, the surrounding tubulo-vesicular spongiome network, and two pores connecting the bladder to the plasma membrane. Dop1p (blue) primarily labels the bladder, and Vma4p (green) exclusively labels the spongiome. Also shown are the oral apparatus (OA) in the anterior end of cell, and both the micronucleus (MIC) and macronucleus (MAC) near the cell midpoint. The vertical rows represent the cytoskeletal tracks called primary meridians, along which the ciliary basal bodies are spaced. For clarity, the cilia themselves are only shown at the cell edges. A, right: As previously reported, Vps8Dp localizes to the CVC as well as to cytosolic puncta. Growing cells expressing Vps8Dp endogenously tagged with mCherry were fixed and stained with DAPI. Images were taken with a Marianas spinning disc confocal microscope. B: *VPS8D* RNAi knockdown strategy via hairpin expression. Cell lines transformed with the hairpin construct, under the control of the inducible MTT1 promoter, are called *VPS8D*-KD. C: *VPS8D* expression is reduced in cells expressing the *VPS8D* hairpin. The accumulation of Vps8Dp-FLAG, in cells expressing the tagged protein from the endogenous locus, was measured by anti-FLAG immunostaining. The labels “-Cd” and “+Cd” indicate the presence or absence of cadmium, which induces *VPS8D* hairpin expression. Images were taken with a Zeiss Axio Observer 7 system. Each image was generated from10 z-sections across the CVC using the Z project tool in FIJI software. D: Quantification of Vps8Dp-FLAG accumulation without/with hairpin-induced knockdown. 10 images for each condition (+Cd and -Cd) were used to quantify the intensity of CVC-localized Vps8Dp immunofluorescence. The data were plotted to a box-and-whiskers plot with statistical analysis via two-tailed *t*-test by GraphPad Software Prism. The boxes display median and interquartile ranges, and the whiskers represent the minimum to maximum value of data. The difference between +Cd and -Cd is significant, *P*-value <0.001. E: *VPS8D* knockdown (*VPS8D*-KD) cells fail to restore normal cell shape after hypoosmotic-induced swelling. DIC images from wild type (WT) and *VPS8D*-KD cells are shown. Images were taken with a Zeiss Axio Observer 7 system. F: *VPS8D* knockdown cells do not survive hypoosmotic stress. Wild type (WT) and *VPS8D*-KD cells were incubated with (+Cd) or without cadmium (-Cd), and then given a hypoosmotic challenge. After 16h, cell viability was measured (see details in Materials and Methods). The data were plotted using GraphPad Software Prism, using a two-tailed *t*-test. Individual data points and the mean±s.d. percentage of survival are shown. The *P*-value for the difference between WT and *VPS8D*-KD is <0.0001. G: Differences in the responses after hypoosmotic shock between wild type and *VPS8D*-KD cells.

### RNAi-mediated knockdown of *VPS8D* results in defects consistent with a role at the osmoregulatory CVC

The role of Vps8Dp at the CVC is not known, but the gene appeared to be essential since we could not recover viable cells when pursuing a gene knockout strategy (Sparvoli et al., 2018). We therefore chose to pursue a gene knockdown approach, by constructing an RNA hairpin corresponding to a part of the VPS8D gene (Fig. 1B) and expressing it in *Tetrahymena* under the control of the cadmium-inducible *MTT1* promoter (Shang et al., 2002, Howard-Till and Yao, 2006). The effectiveness of such hairpins in *T. thermophila* can be difficult to assess at the level of the targeted transcripts (Howard-Till and Yao, 2006). To circumvent this issue, we directly measured protein levels by generating a cell line in which Vps8Dp was endogenously tagged with the FLAG epitope, and the *VPS8D* hairpin was inducibly expressed from the *RPL29* gene locus (Yao and Yao, 1991) (Fig. 1C). Induction of the hairpin expression led to a significant decrease in the level of Vps8Dp-FLAG, as measured by immunofluorescence (Fig. 1D). Importantly, after hypoosmotic shock, the knockdown cells displayed morphological changes consistent with a defect in maintaining osmotic homeostasis, losing their teardrop shape and becoming roughly spherical (Fig. 1E) Equivalently-treated wild type cells could quickly recover their normal shape (Fig. 1E). Similarly, while the wildtype cells retained full viability after the osmotic challenge, the *VPS8D* knockdown cells died shortly thereafter (Fig. 1F). The differences in the cell responses after hypoosmotic shock between wild type and *VPS8D*-KD cells are diagrammed in Fig. 1G. Under these stress conditions, the activity of the CVC appears critical not to prevent osmotic swelling but to allow recovery after such events.

### *VPS8D* knockdown affects features of both the bladder and spongiome

Conventional CORVET in *S. cerevisiae* acts as a determinant in the maturation of endosomal compartments (Peplowska et al., 2007). To ask whether knockdown of *T. thermophila VPS8D* led to changes in the CVC, we generated cells expressing the *VPS8D* hairpin in combination with endogenously tagged copies of either Dop1p, which most strongly labels the bladder, or tagged Vma4p which decorates the spongiome (Cheng et al., 2023).

Following 5 hours of hairpin induction, Dop1p-mNeon still shows clear localization to the CV bladder that is qualitatively similar to that in non-induced or wildtype cells, and the bladder profiles are also similar (Fig. 2A, compare left and right panels). However, by 8 hours of hairpin induction the intensity of Dop1p bladder steady state labeling was significantly decreased (Fig. 2B and S2). This change in the association between Dop1p and the bladder membrane upon *VPS8D* knockdown could also be seen at the level of dynamics. We previously used fluorescence recovery after photobleaching (FRAP) to discover that Dop1p at the bladder exchanges with a cytosolic pool (Cheng et al., 2023), as shown in Figure 2C and 2D. That exchange was inhibited in *VPS8D* knockdown cells, so that the rate of fluorescence recovery at the bleached bladder was decreased (Fig. 2E, 2F and 2G). Lastly, the period between contractions was lengthened in these *VPS8D* knockdown cells (Fig. 2H). The mechanisms controlling bladder contraction are not known, but one possibility is that bladder refilling is less efficient in the mutant. Intriguingly, *VPS8D* hairpin induction had a clear effect on the spongiome morphology. In particular, the volume occupied by Vma4p-decorated tubules showed a marked decrease within 5 hours of *VPS8D*-hairpin induction (Fig. 2I, 2J and S3), suggesting that spongiome architecture depends more acutely than bladder structure on the activity of Vps8Dp. Interestingly, these rapid changes in spongiome structure upon *VPS8D* knockdown are similar to those seen after cells are exposed to brief hypoosmotic challenge (Fig. S4).

**Figure 2:**
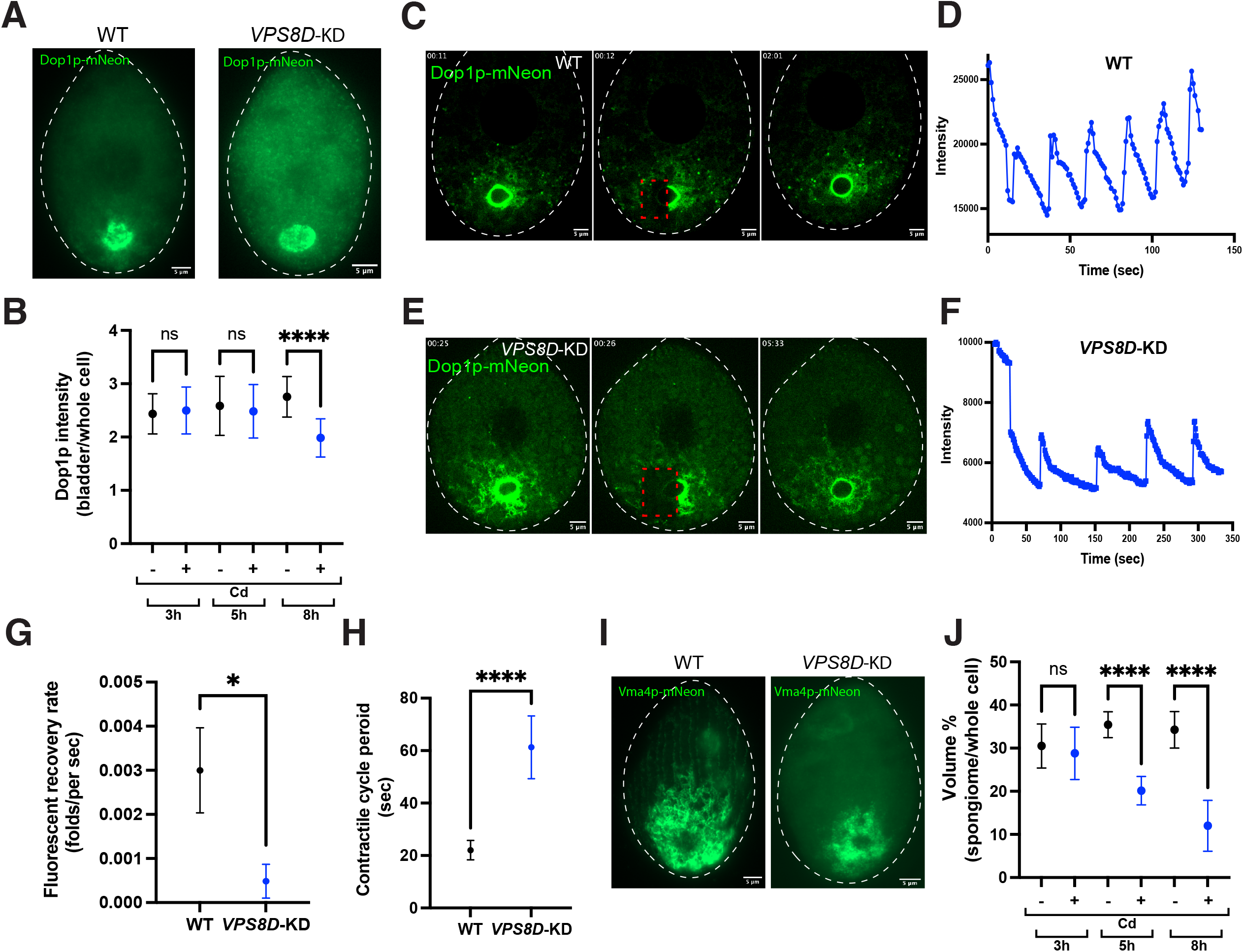
*VPS8D* is required for features of the bladder and spongiome. A: Dop1p localization becomes more dispersed upon *VPS8D* knockdown. Growing cells (WT or *VPS8D-*KD) expressing Dop1p-mNeon at the *DOP1* locus were imaged to assess Dop1p localization. WT (left panel); induced *VPS8D* knockdown (right panel). Images were taken with a Zeiss Axio Observer 7 system. B: Dop1p localization to the bladder decreases upon *VPS8D* knockdown. *VPS8D*-KD cells expressing Dop1p-mNeon were incubated without (-Cd) or with (+Cd) cadmium for 3,5 or 8 hours and then fixed for imaging. 20 images for each sample were analyzed to quantify Dop1p fluorescence at the bladder and for the whole cell (Fig. S2). The data were plotted using GraphPad Software Prism and analyzed using a two-tailed *t*-test. The ratios of the mean±s.d. intensities of Dop1p fluorescence at the bladders relative to the whole cells from each sample are shown. The difference between +Cd and -Cd after 8-hour treatment is significant, *P*-value <0.0001. C-G: Dop1p exchange kinetics are slowed upon *VPS8D* knockdown. C. Wild type cell expressing Dop1p-mNeon analyzed by FRAP. Growing cells were immobilized and live imaged with a Marianas spinning disc confocal microscope with a FRAP tool. Three images were extracted from a video (Movie 1) showing a photobleaching event and recovery. Left panel: cell at 11s, immediately before photobleaching. Middle panel: cell at t=12s immediately after photobleaching a region containing part of the CV bladder. The bleached area appears as a dark square area outlined with a red dotted box. Right panel: cell at t=2 min 1s. D: The recovery after photobleaching data were analyzed by FIJI image process software (see details in Materials and Methods). The fluorescence intensity in the photobleached area along with the time was plotted using GraphPad Software Prism. E: *VPS8D* knockdown cell expressing Dop1p-mNeon analyzed by FRAP. Three images were extracted from a video (Movie 2) showing a photobleaching event and recovery, with panels as in Fig. 2C but with the bleaching event at t=0:26 and recovery at t=5:33. F: Recovery after photobleaching, plotted as in Fig. 2D. G: Dop1p recovery after photobleaching is slowed in *VPS8D* knockdown cell. Three FRAP experiments obtained from wild type and *VPS8D* knockdown cells were analyzed and calculated to obtain the mean±s.d. recovery rate after photobleaching. The data were plotted using GraphPad Software Prism and using a two-tailed *t*-test. The difference between wild type and *VPS8D* knockdown is significant, *P*-value <0.05. H: Contractile cycle period is increased in *VPS8D* knockdown cells. The contractile cycle period of wild type cell and *VPS8D* knockdown cell were measured. 50 cycles from wild type cells and 22 cycles from *VPS8D* knockdown cells were measured to obtain the contractile cycle period and the data were plotted using GraphPad Software Prism and using a two-tailed *t*-test. The difference between wild type and *VPS8D* knockdown is significant, *P*-value <0.0001. I: Vma4p localization in WT and *VPS8D* knockdown cells. Growing cells expressing Vma4p-mNeon were imaged to show Vma4p localization in WT (left panel) and *VPS8D* knockdown cells (right panel). Images were taken with a Zeiss Axio Observer 7 system. J: The spongiome area is reduced in *VPS8D* knockdown cell. *VPS8D*-KD growing cells expressing Vma4p-mNeon were treated with cadmium (+Cd) for 3,5 and 8 hours or without cadmium (-Cd) and then were fixed for imaging. 20 images for each sample were analyzed to quantify the area of the Vma4p-labelled spongiome and the whole cell (Fig. S3). The data were plotted using GraphPad Software Prism and using a two-tailed *t*-test. The ratio of the mean±s.d. volume of spongiome to whole cell from each sample is shown. The difference between +Cd and -Cd at 5- and 8-hours treatment point are significant, *P*-value <0.0001.

### Prolonged induction of *VPS8D* hairpin expression leads to loss of CVC structures

Following longer periods of *VPS8D*-hairpin induction, the CVC and associated markers underwent more dramatic changes. Strikingly, many cells viewed with DIC optics lacked any visible contractile bladders, strongly suggesting that the bladders are lost during sustained *VPS8D* knockdown. Not surprisingly, all CVC proteins in such cells showed a marked change in their distribution compared to in wildtype. Dop1p no longer localizes to a recognizable bladder but is instead dispersed throughout the cytoplasm including in many small puncta (Fig. 3A). Vma4p is also dispersed throughout the cell, visible as diffuse fluorescence but also present in large, potentially multi-vesicular structures (Fig. 3B). A second protein that localizes to the spongiome in wildtype cells, Scr7p, is also delocalized after extended *VPS8D*-hairpin induction. The delocalization of these two spongiome markers is likely to reflect the loss of the spongiome, in parallel to the loss of the bladder.

**Figure 3:**
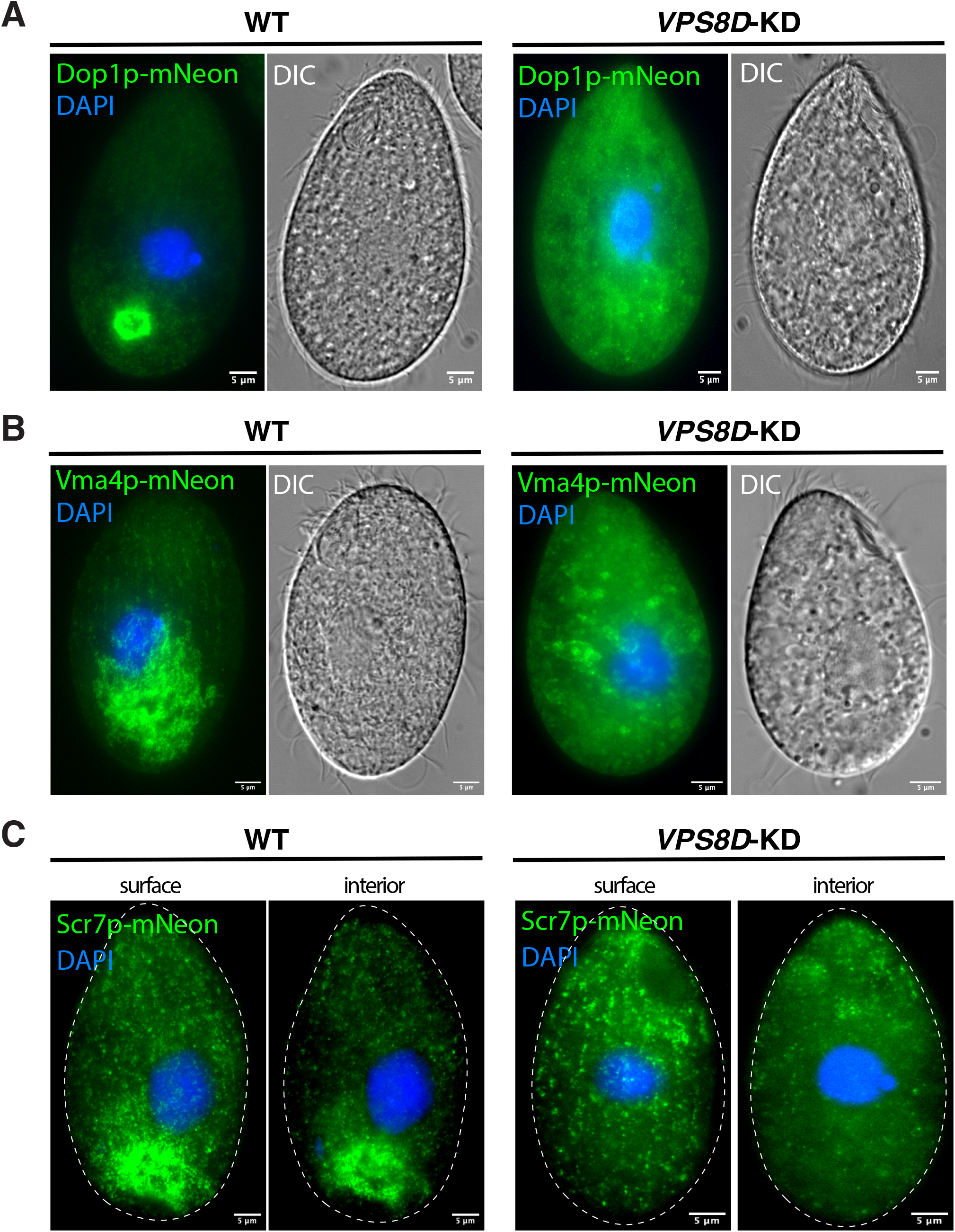
Prolonged *VPS8D* hairpin induction leads to cytoplasmic dispersal of Dop1p, Vma4p and Scr7p. A: Severe mislocalization of Dop1p upon *VPS8D* knockdown. Growing cells expressing *DOP1*-mNeon were fixed and the nuclei stained with DAPI. DIC and fluorescent images of the wild type and *VPS8D* knockdown cells are shown in the left and right panels, respectively. B: Severe mislocalization of Vma4p upon *VPS8D* knockdown. Growing cells expressing *VMA4*-mNeon were fixed and the nuclei stained with DAPI. DIC and fluorescent images of the wild type and *VPS8D* knockdown cells are shown in the left and right panels, respectively. C: Severe mislocalization of Scr7p upon *VPS8D* knockdown. Growing cells expressing *SCR7*-mNeon were fixed and the nuclei stained with DAPI. Fluorescent images of wild type and *VPS8D* knockdown cells are shown in the left and right halves, respectively, with focal planes selected from Z-stacks to illustrate the cell surface (left) and cell interior (right). Images were taken with a Zeiss Axio Observer 7 system.

Taken together, these results strongly suggest that the maintenance of multiple compartments of the CVC depend on Vps8Dp. The spongiome may be more acutely dependent on Vps8Dp-assisted membrane trafficking than the bladder.

### *VPS8D* knockdown results in a proliferation of large vesicles or vacuoles bearing CV markers

If Vps8Dp as part of a CORVET complex is active in promoting fusion between membranes, then the disappearance of CVC structures upon *VPS8D* knockdown may be due to a fusion defect between intermediates in CVC formation. To explore this idea, we analyzed cells that showed intermediate CVC phenotypes after induction of the *VPS8D* hairpin. In these experiments, cells were treated with low concentrations of cadmium and for short time periods. In such samples, it was easy to find cells displaying CVC features never seen in wildtype cells, with examples in Figures 4A (wildtype) vs the knockdown cells (Fig. 4B and 4C). The cell shown in panel B possesses what appear to be two adjacent CV bladders. These persist for tens of seconds before one of them contracts rapidly (t = 28sec), which is then followed by contraction of the second bladder at 29 sec. This asynchronicity implies that the bladders are linked with two different plasma membrane pores. Over the next 20 seconds, just one of the bladders appears to refill. In panel C, the cell contains at least seven medium sized-to-large Dop1p-labeled vesicles. Over the course of the 15 seconds in the video, one of the two largest vesicles contracts, followed shortly thereafter by the second. Within 5-6 seconds, both show signs of refilling and also appear tightly associated with smaller labeled vesicles at their periphery. The Dop1p-labeled relatively immobile vesicles in this cell are spread over a large volume of the cell posterior, so that most cannot be closely associated with CV pores. The presence in a large subset of the knockdown cells of heterogeneous Dop1p-labeled compartments, which include multiple contractile bladders, may be consistent with the idea that maintenance of the CVC in wildtype cells requires Vps8Dp-dependent membrane fusion.

**Figure 4:**
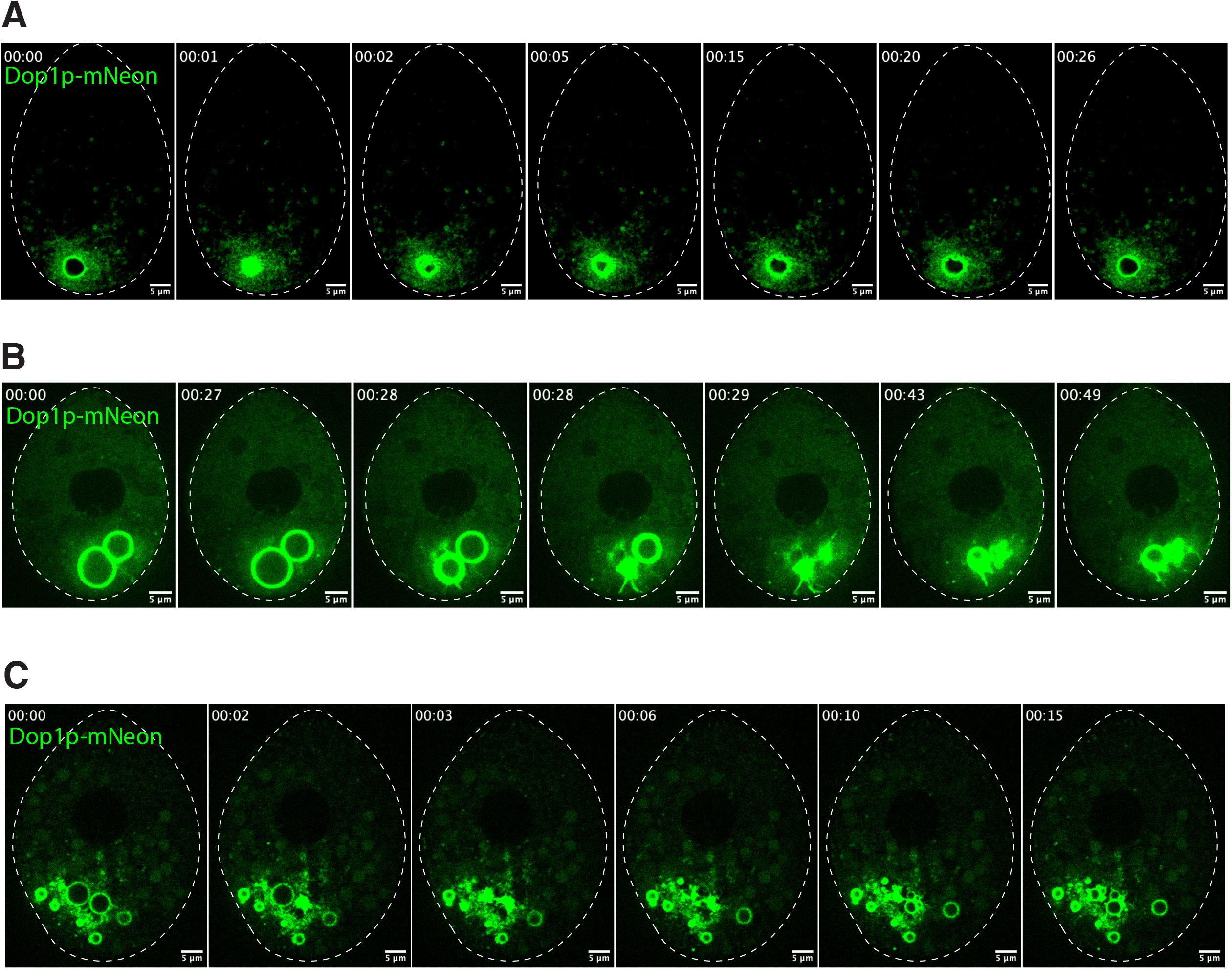
Live imaging of Dop1p reveals altered CVC morphology upon *VPS8D* knockdown. A: Live imaging of the CVC contractile cycle in a wildtype cell expressing Dop1p-mNeon. The focal plane represents a cross-sectional view of the CVC, with 0.5 sec frame intervals. Seven successive images extracted from a video (Movie 3) are shown. Cells from growing cultures were immobilized using CyGEL, as described in Materials and Methods. Videos were captured using a Marianas spinning disc confocal microscope. B: Live imaging of the CVC in *VPS8D* knockdown cell expressing Dop1p-mNeon. The cell selected contains two large bladders that undergo asynchronous contractions. Imaging conditions were the same as in Fig. 4A. The seven successive time-lapse images were extracted from Movie 4. C: Live imaging of the CVC in *VPS8D* knockdown cell expressing Dop1p-mNeon. The cell selected contains multiple Dop1p-labeled vesicles dispersed in the cell posterior, concentrated in the region where the single stereotypical CVC bladder is typically located. Two of them contract over the course of the video. Imaging conditions were the same as in Fig. 4A, with 1 sec frame intervals. The six successive time-lapse images were extracted from Movie 5.

### Vps8Dp is concentrated at junctions between CV-related membranes

In budding yeast, the HOPS complex concentrates at the junctions between membranes destined for fusion during vacuole maintenance (Wang et al., 2003).To see whether Vps8Dp in *Tetrahymena* behaves similarly, we took advantage of the observation that many cells exposed to hypoosmotic stress showed fragmented CVC structures. As shown in Figure 5A, such cells contain multiple Dop1p-labeled vacuole-like compartments, concentrated in the cell posterior and undergoing frequent fusion to form larger compartments, in addition to what appears to be the central contractile bladder. Dop1p itself appears to be irregularly distributed, often as puncta, around the periphery of the labeled compartments. Interestingly, the CVC similarly fragments into multiple Dop1p-decorated large vacuole-like compartments under conditions of <u>hyper</u>osmotic stress, though with the difference that no contractile central bladder is visible (Fig. 5B).

**Figure 5:**
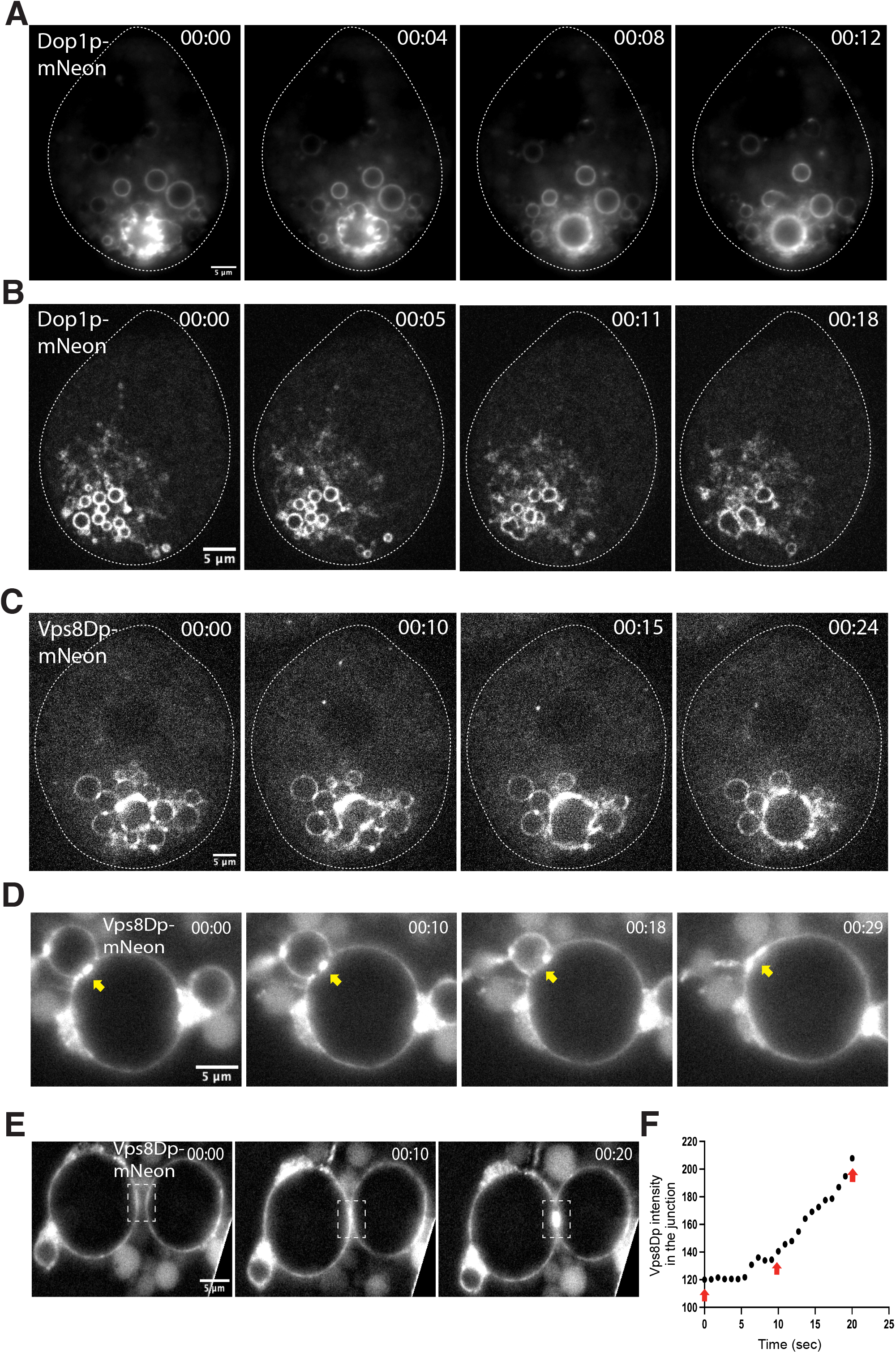
Vps8Dp concentrates at junctions between adjacent vacuoles. A: Live imaging of numerous large Dop1p-labeled vacuoles in cells expressing Dop1p-mNeon, following <u>hypo</u>-osmotic shock. The cell shown was imaged to capture a cross-sectional view, in a time-lapse video with 4 sec frame intervals (Movie 6). Four successive images extracted from the video are shown. Growing cell cultures were hypo-osmotically challenged with 10 mM Tris-HCl buffer, pH 7.4 and immediately immobilized for imaging using CyGEL, as described in Materials and Methods. Videos were captured using a Zeiss Axio Observer 7 system. B: Live imaging of the numerous Dop1p-labeled vacuoles following <u>hyper</u>-osmotic shock in cells expressing Dop1p-mNeon. The cell shown was imaged to capture a cross-sectional view, in a time-lapse video with 0.15 sec frame intervals (Movie 7). Four successive images extracted from the video are shown. Growing cell cultures were hyper-osmotically challenged with 10 mM sorbitol and immediately immobilized for imaging using CyGEL, as described in Materials and Methods. Videos were captured using a Marianas spinning disc confocal microscope. C: Live imaging of numerous large Vps8Dp-labeled vacuoles following <u>hypo</u>-osmotic shock in cells expressing Vps8Dp-mNeon. The cell shown was imaged to capture a cross-sectional view, in a time-lapse video with 0.5 sec frame intervals (Movie 8). Four successive images extracted from the video are shown. Cells were treated as described in Fig. 5A. Videos were captured using a Marianas spinning disc confocal microscope. D-E: Imaging of extracellular Vps8Dp-labeled vacuoles that leaked from ruptured cells expressing Vps8Dp-mNeon. D. Vps8Dp localizes densely at contact site between adjacent vacuoles, which in some cases are sites of subsequent membrane fusion (arrowhead). Four successive images extracted from Movie 9 are shown. E: Vps8Dp re-distributes at the time that two vacuoles come into contact. The two large vacuoles shown were separated by ∼1µ at the beginning of the video (t=0) (Movie 10). When they moved into contact starting at t=10s, Vps8Dp accumulated at the junction (boxed). A second example of this phenomenon is shown in Movie 11. F. Plot of fluorescence intensity change with time for the boxed area shown in Fig. 5E, in Movie 10. The data were plotted using GraphPad Software Prism. The three red arrows correspond to the images shown in panel E.

Vacuole-like compartments formed under hypoosmotic conditions are also decorated with Vps8Dp (Fig. 5C). The Dop1p-labeled and Vps8Dp-labeled vacuoles appear to be the same structures, since there is strong co-localization of the two proteins at such vacuoles in cells expressing both Dop1p-mNeon and Vps8Dp-mCherry (Fig. S5). However, a notable difference between the Dop1p and Vps8Dp distributions is that in many cases the latter appears strikingly concentrated in regions between adjoining vacuoles, where the membranes come into close contact (Fig. 5C).

The concentration of Vps8Dp-mNeon between adjoining vacuoles/vesicles could be seen even more strikingly in some osmotically-stressed samples in which cells occasionally suffered localized plasma membrane rupture under the pressure from the microscope cover slip. These breaks allowed leakage of the large Vps8Dp-labeled vesicles into the surrounding buffer. In such samples, Vps8Dp could be seen highly concentrated at junctures between the membranes, consistent with a role in tethering and fusion (Fig. 5D). Moreover, in some cases we could capture membrane fusion occurring at Vps8Dp-enriched junctions (Fig. 5D, arrowheads). Interestingly, we also captured episodes in which two of the leaked vesicles shifted closer to one another, and Vps8Dp accumulated rapidly at their meeting site just when they came into contact (Fig. 5E and 5F).

## Discussion

Genes encoding CORVET subunits underwent a marked expansion in the lineage leading to Tetrahymena, leading to the six distinct complexes in *T. thermophila* called 8A-8F based on their unique Vps8 subunit (Klinger et al., 2013, Sparvoli et al., 2018, Sparvoli et al., 2020). These complexes are distinct from one another in their localization and their subunit makeup (Sparvoli et al., 2020). They are accordingly likely to be individually specialized for distinct functions. We confirmed this previously for the 8A complex, which we found to be essential for the formation of lysosome-related organelles called mucocysts (Sparvoli et al., 2018).

In this manuscript we report analysis of the Vps8 subunit of the 8D CORVET. Vps8Dp was previously shown to be strongly associated with the osmoregulatory CVC (Sparvoli et al., 2020, Cheng et al., 2023). Because the gene was found to be essential for cell viability and therefore difficult to analyze via gene knockout (Sparvoli et al., 2018), in this manuscript we instead used a knockdown approach based on induced expression of a gene-specific RNA hairpin (Howard-Till and Yao, 2006). One striking finding was that even relatively minor depletion of Vps8Dp produced cells that were highly sensitive to osmotic shock (Fig. 1). More informatively, following hairpin induction the cells showed time-dependent changes in the morphology and functioning of the CVC, affecting both the spongiome and bladder.

Within the cohorts of knockdown cells in each experiment there was significant phenotypic heterogeneity, making it difficult to rigorously establish the order in which different defects were manifest. Nonetheless, our results suggest that spongiome structure may be more sensitive to *VPS8D* dosage compared with the bladder structure, since at early timepoints following hairpin induction only the spongiome was reduced in size relative to the wildtype, a response that became more pronounced at longer time points (Fig. 2J). Interestingly, this shrinkage of the spongiome is very similar to the response seen when cells are briefly exposed to hypoosmotic shock (Cheng et al., 2023). The bladder size in contrast did not detectibly change, but contractions grew less frequent (Fig. 2G). Since the bladder fills with water transported from the spongiome, the slowed contraction rate may reflect less efficient filling from a partially disabled water-collecting reticulum. The bladder in knockdown cells at these early time points also showed reduced labeling by Dop1p, with a corresponding increase in diffuse Dop1p signal in the cytoplasm (Fig. 2A and 2B). The mechanistic role of Dop1p is not yet understood, but in its absence cells show only infrequent contraction of an enormously expanded bladder (Cheng et al., 2016, Cheng et al., 2023).

At later time points following hairpin induction, neither the bladder nor spongiome was detectable, indicating that *VPS8D* is essential for the maintenance of this organelle (Fig. 3). Proteins that in wildtype cells localized strongly to the CVC were instead found in dispersed cytoplasmic puncta. This process may be reversible, based on our observation that some cells in knockdown cultures recover when hairpin expression is de-induced. Studying the mechanisms underlying such recovery may help to shed light on trafficking pathways involved in CVC biosynthesis, which are poorly understood in any lineage and particularly in Ciliates. The disappearance of both bladder and spongiome compartments after prolonged *VPS8D* knockdown is similar to a phenotype reported in *D. discoideum* that is linked to a protein in the BEACH family, called LvsA (Gerald et al., 2002). Interestingly, pulldown experiments in the Apicomplexan *Toxoplasma gondii* suggest physical interactions between CORVET/HOPS and a BEACH-domain containing protein, but in the context of secretory organelle biogenesis (Morlon-Guyot et al., 2018).

The established role of CORVETs, based on studies in yeast and animals, is in tethering and fusion of membrane compartments (Spang, 2016, van der Beek et al., 2019). Our imaging data are consistent with such a role for the Vps8Dp-containing CORVET at the CVC. For our imaging, we exploited conditions in which the CVC bladder, rather than existing as a single large vacuole, is replaced by a set of smaller vacuoles that can still be recognized due to their labeling by Dop1p as well as Vps8Dp. We found that Dop1p has a relatively uniform distribution at the periphery of such vacuoles, while in contrast Vps8Dp is concentrated at sites where neighboring vacuoles come into close contact (Fig. 5). In some cases, these were seen to be sites of subsequent fusion. A similar phenomenon has been reported of specific protein and lipid accumulation at the interface between vacuoles undergoing homotypic fusion in yeast, termed the vertex ring domain, and it includes the HOPS complex (Wang et al., 2003, Fratti et al., 2004). Interestingly, we witnessed the same highly polarized Vps8Dp distribution on small Vps8Dp-decorated vacuoles that were extruded through breaches in the plasma membrane of perforated cells. In one case, we witnessed rapid Vps8Dp redistribution at the perimeters of two initially widely-spaced vacuoles when they came into contact, leading to accumulation at the junction (Fig. 5E). This may be explained if the 8D-CORVET complexes on each surface are bound to integral membrane proteins undergoing rapid 2-dimensional diffusion, but any complex that simultaneously engages with integral membrane proteins on an adjacent membrane diffuses more slowly. Based on analysis of the vertex ring complex in yeast, this partitioning is also likely to involve lipid subdomains (Fratti et al., 2004).

Our images from osmotically-stressed cells are consistent with a role for Vps8Dp in homotypic tethering and fusion at the CVC, and therefore support the paradigm for CORVET and HOPS established in other eukaryotes. However, no Vps8Dp-labeled vertex ring domains are apparent in cells grown under non-stress conditions, and moreover there is no step clearly involving homotypic fusion in the *Tetrahymena* contractile cycle (Cheng et al., 2023). In Paramecium, electrophysiological data support a model in which the junctions between the bladder and emanating spongiome arms undergo fission and subsequent fusion with each contractile cycle (Tominaga et al., 1998b). Whether similar fission/fusion cycles occur in Tetrahymena, where the structure of the bladder/spongiome junction appears quite different, is unknown (Elliott and Bak, 1964). The role of Vps8Dp is unlikely to be restricted to either homotypic fusion between bladder-derived vesicles or spongiome-bladder fusion, since Vps8Dp localizes to puncta throughout the spongiome (Cheng et al., 2023). Spongiome tubules show dynamic extension and branching (Cheng et al., 2023), and one possibility is that the Vps8Dp-containing CORVET complex facilitates homotypic fusion between branches in a way that is required to maintain a cohesive reticulum. The retraction of the spongiome at early time points following *VPS8D* knockdown may be consistent with this scenario.

However, another broad possibility is that Vps8Dp has activities independent of a conventional CORVET complex, as a monomer or a smaller sub-complex, for which examples exist in other eukaryotes for CORVET and HOPS (Asensio et al., 2013, Lorincz et al., 2016). In pulldowns, Vps8Dp is associated with the 5 other expected subunits of CORVET (Sparvoli et al., 2020). However, some details suggest it may be an outlier among the *Tetrahymena* CORVET complexes. First, it is the most biochemically differentiated, including unique subunit variants for Vps8, Vps16, Vps18, and Vps33 (Sparvoli et al., 2020). Secondly, Vps8Dp is roughly 200 amino acids longer than the other Vps8 paralogs in Tetrahymena, with multiple insertions suggesting the potential for additional interactions and activities. Lastly, Vps8Dp in pulldowns appeared by silver staining to be significantly more abundant than the other subunits with which it associates; significantly, no similar imbalance was present for the Vps8 subunits of the other five CORVETs (Sparvoli et al., 2020). Since the affinity tag for these pulldowns was attached to Vps8Dp, the greater relative abundance of Vps8Dp could be explained if that particular CORVET were comparatively unstable. Alternatively or in addition, it could reflect a super-stoichiometric pool of Vps8Dp in cells. On this basis, we speculate that Vps8Dp could potentially be playing distinct roles at the bladder vs. spongiome, as part of different complexes.

## Materials and Methods

### Cell strains and culture conditions

*Tetrahymena thermophila* strains used in this work are listed in Table S1 with culture conditions as previously described (Gorovsky et al., 1975). Cells were grown in SPP medium (2% proteose peptone (GIBCO, 211684), 0.1% yeast extract (BD, 212750), 0.2% dextrose (ACROS, 41095-5000) and 0.003% EDTA, ferric-sodium salt) supplemented with 100 μg ml^-1^ Normocin^TM^ (InvivoGen), an antimicrobial reagent. Cells were grown at 30°C with agitation at 100 rpm. Cell densities were measured using a spectrophotometer (Thermo Spectronic Unicam), where OD_550_ = 1 corresponds to ∼ 1 *×* 10^6^ cells ml ^-1^ (David L. Spector, 1998) or a Z1 Coulter Counter (Beckman Coulter Inc., Indianapolis, Indiana). Cultures were generally used during the log phase of cell growth (cell density 2-4 × 10^5^ cells ml^-1^). Reagents were from Sigma-Aldrich unless otherwise noted.

### Biolistic Transformation

50 ml *Tetrahymena* cultures were grown to 500,000 cells ml^-1^ and starved for 10–16 h in 10 mM Tris-HCl buffer, pH 7.4, at 30°C with agitation. Biolistic transformations were performed as described previously (Sparvoli et al., 2018, Cassidy-Hanley et al., 1997). Transformants were identified after 3 days selection in paromomycin sulfate (120 μg ml^-1^ with 1 μg ml^-1^ of CdCl_2_), and then serially passaged 5×/week for ∼4 weeks in decreasing concentrations of CdCl_2_ and increasing concentrations of paromomycin sulfate.

### Generation of *VPS8D* knockdown strains

To generate *VPS8D* knockdown strains, a construct was designed as shown in Fig. 1B, by first PCR amplifying ∼500 bp of genomic *VPS8D* gene coding sequence with two sets of primers, one set engineered with ApaI and XhoI restriction sites and another set with PmeI and SmaI restriction sites. These two fragments were cloned into PCRII-I3 so that they were inverted relative to one another. The rDNA backbone vector pIBF (Howard-Till and Yao, 2006) was digested with with ApaI and PmeI to release the BFP fragment. In parallel, the *VPS8D* hairpin cassette was excised from the PCRII-I3 backbone with ApaI and PmeI, and the ∼1kb insert then gel purified and ligated into the site of the excised BFP fragment in the pIBF backbone. Approximately 20µg of this plasmid was used per biolistic transformation. In addition to the rDNA-based *VPS8D* hairpin strains, we also engineered a construct to target the *VPS8D* hairpin cassette to the macronuclear *RPL29* locus. In this case, the*VPS8D* hairpin cassette was cloned into pBSICC-rpl29CyR (Yao and Yao, 1991) for biolistic transformation followed by selection for cycloheximide resistance. In both *VPS8D* hairpin constructs, hairpin expression is under the control of the cadmium-inducible *MTT1* promoter (Shang et al., 2002).

### *Assaying VPS8D* knockdown phenotypes

To measure survival rates after hypoosmotic shock, wildtype and *VPS8D*-KD strains (3 of each, with independent isolates of the mutant strain) were grown to log phase in 10 ml SPP. Cell densities were measured and equalized, and 0.5ml aliquots were diluted 1:100 in 50 ml SPP medium with or without 1 µg/ml CdCl_2_. Cells were grown for 16 hours at 30°C with agitation. Culture densities were measured, and then cells were pelleted by centrifugation in a clinical centrifuge for 1 minute and resuspended gently in 10 mM Tris-HCl buffer, pH 7.4. After an overnight incubation, culture densities were once again measured to calculate survival rates. For experiments monitoring changes in CVC markers, the *VPS8D*-KD strains endogenously expressing Dop1p-mNeon, Vma4p-mNeon or Scr7p-mNeon were grown as above in non-inducing or inducing conditions. Cells were withdrawn and fixed for imaging after 3, 5 and 8 (Fig. 2) or 16h (Fig. 3).

### Acute osmotic challenge

Cells expressing tagged proteins were grown to log growth phase, then pelleted by centrifugation in a clinical centrifuge for 1 min and gently resuspended in 22°C 10 mM Tris-HCl buffer, pH 7.4 (hypoosmotic) or 10 mM sorbitol (hyperosmotic). The cells were then immediately immobilized for live imaging.

### Endogenous expression of FLAG-tagged Vps8Dp and mNeon-tagged genes in ***VPS8D* knockdown strains**

pVPS8D-FLAG-ZZ-Neo4, pDOP1-2mNeon-6myc-Neo4, pVMA4-2mNeon-6myc-Neo4 and pSCR7-2mNeon-6myc-Neo4 (Cheng et al., 2023) were introduced by biolistic transformation into strains in which the *VPS8D* hairpin expression cassette was already integrated at the *RPL29* locus, as described above.

### Cell immobilization for live imaging

We used two methods to immobilize cells for live imaging. For both methods, we prepared a single slide at a time and viewed it immediately. The first method relied on the pressure exerted on small volume samples under a cover slip. Cells were first concentrated by centrifugation to 2-5 x 10^6^ cells ml^-1^. 6 μl of sample was applied to the slide, and immediately overlayed with a 22 × 22 mm cover slip. The resulting pressure frequently results in cell immobilization, which could be verified by monitoring ciliary beating on the cell surface at the same time that cell viability could be monitored by observing the periodic contraction of the contractile vacuole, both in the DIC channel. For the second method, cells were pelleted and resuspended in a thermoreversible gel on ice, whose viscosity increases as it warms to room temperature. Here, cells at the same high density were mixed well with CyGEL^TM^ (ab109204, ABCAM) (the mix ratio optimized for each experiment) and immediately mounted with a cover slip. This approach results in many cells being effectively immobilized in the thin gel. For both methods, we carefully watched for proliferation of large cytosolic vesicles. Growing cells have large food vacuoles derived from the oral apparatus. No new food vacuoles form in immobilized cells, so any substantial increase in the number of such vesicles is likely to represent autophagosome formation as part of a stress response. We rejected any cells showing such an increase.

### Cell fixation for imaging

Cells were fixed in a final concentration of 4% paraformaldehyde in 10 mM Tris-HCl buffer, pH 7.4 (stock: 16% paraformaldehyde solution in distilled water, EM grade, 15710, Electron Microscopy Science) for 10-30 minutes at room temperature. They were then washed repeatedly in PBS to reduce the background autofluorescence, before staining with DAPI (4′,6-diamidino-2-phenylindole) at final concentration 50 ng ml^-1^ for 10 minutes for nuclear staining, and/or staining with antibodies for immunofluorescence. Images were captured by using a Carl Zeiss Microscope stand Axio Observer 7 system or Marianas Yokogawa-type spinning disc inverted confocal microscope.

### Fluorescence Recovery after Photobleaching (FRAP)

Cells were immobilized as described above. FRAP experiments were performed using a Marianas Yokogawa-type spinning disc inverted confocal microscope with a 100x/NA1.45 oil (Alpha Plan-Fluar) objective. The system featured fast shutter speeds and channel switching for high-speed imaging and a vector high-speed point scanner for bleaching, controlled using Slidebook software. For each experiment, we first identified a well-immobilized cell and verified the fluorescence signal with a ∼100-500 ms exposure. We then used the time-lapse mode to capture images with the fastest camera speed for a pre-photobleaching record and then used the screen selection tool and real-time imaging to draw a ROI (region of interest) for photobleaching.

### Image analysis

The image processing package FIJI was used for image processing and analysis (Schindelin et al., 2012). Processing of raw images included: image cropping and rotation; bleach correction; background subtraction; adjustment of brightness/contrast; color switching; selection of images from z-stacks; intensity threshold adjustments; measurements of parameters including areas, mean gray values, and integrated densities.

To quantify Dop1p-mNeon fluorescence intensity in the CV bladder vs whole cell, all images were taken using the same image capturing settings. The raw images were used to select the ROIs of the area of Dop1p-labeled bladder and whole cell. The mean fluorescence intensities of these ROIs were then measured. To quantify Vma4p-mNeon localized spongiome distribution, the raw images were used to measure the area of Vma4p-labeled spongiome and whole cell. For FRAP analysis, the time-lapse images were separately adjusted by bleach correction for the pre-photobleaching series and after-photobleaching series. The intensity thresholds were adjusted for background subtraction, and intensity changes over time were measured in ROIs. Intensities were plotted using GraphPad Software Prism.

### Immunofluorescence analysis

Cells (2×10^5^) were fixed with 4% paraformaldehyde for 30 minutes at room temperature, and immunolabeled as previously described (Briguglio et al., 2013, Bowman and Turkewitz, 2001). Briefly, cells were first incubated for blocking with 5% BSA (bovine serum albumin, BP1600-1 Fisher Scientific) in PBT (0.3% Triton X-100 (ACROS) in PBS) for one hour at room temperature and then incubated with the primary antibodies in the same blocking buffer. Anti-FLAG M2 antibody staining was at 1:1000 (Monoclonal ANTI-FLAG M2 antibody, Sigma F1804) for overnight at 4°C, followed by three washes in 0.3% Triton X-100 in PBS. The secondary antibodies were Alexa Fluor 488-conjugated anti-mouse IgG, incubated for one hour at room temperature (1:1000, Alexa Fluor 488 Goat anti-Mouse IgG, Invitrogen A-11001), and similarly washed. Cell nuclei were stained with 4′,6-diamidino-2-phenylindole (DAPI) (50 ng ml^-1^). Sample was mounted on microscopic slides for imaging.

## Acknowledgements

At the University of Chicago, we thank Vytas Bindokas and Christine Labno for help at the Integrated Light Microscopy Core Facility. In the Turkewitz laboratory, we thank Patrick Jiang and Maya Waarts for helpful discussions.

## Reference

Allen, R. D. 2000. The contractile vacuole and its membrane dynamics. Bioessays, 22, 1035–42.

Allen, R. D. & Naitoh, Y. 2002. Osmoregulation and contractile vacuoles of protozoa. Int Rev Cytol, 215, 351–94.

Asensio, C. S., Sirkis, D. W., Maas, J. W., Jr., Egami, K., To, T. L., Brodsky, F. M., Shu, X., Cheng, Y. & Edwards, R. H. 2013. Self-assembly of VPS41 promotes sorting required for biogenesis of the regulated secretory pathway. Dev Cell, 27, 425–37.

Baker, R. W., Jeffrey, P. D., Zick, M., Phillips, B. P., Wickner, W. T. & Hughson, F. M. 2015. A direct role for the Sec1/Munc18-family protein Vps33 as a template for Snare assembly. Science, 349, 1111–4.

Bowman, G. R. & Turkewitz, A. P. 2001. Analysis of a mutant exhibiting conditional sorting to dense core secretory granules in Tetrahymena thermophila. Genetics, 159, 1605–16.

Bright, L. J., Kambesis, N., Nelson, S. B., Jeong, B. & Turkewitz, A. P. 2010. Comprehensive analysis reveals dynamic and evolutionary plasticity of Rab GTPases and membrane traffic in Tetrahymena thermophila. PLos Genet, 6, e1001155.

Briguglio, J. S., Kumar, S. & Turkewitz, A. P. 2013. Lysosomal sorting receptors are essential for secretory granule biogenesis in Tetrahymena. J Cell Biol, 203, 537–50.

Cassidy-Hanley, D., Bowen, J., Lee, J. H., Cole, E., Verplank, L. A., Gaertig, J., Gorovsky, M. A. & Bruns, P. J. 1997. Germline and somatic transformation of mating Tetrahymena thermophila by particle bombardment. Genetics, 146, 135–47.

Cheng, C. Y., Romero, D. P., Zoltner, M., Yao, M. C. & Turkewitz, A. P. 2023. Structure and dynamics of the contractile vacuole complex in Tetrahymena thermophila. J Cell Sci.

Cheng, C. Y., Young, J. M., Lin, C. Y. G., Chao, J. L., Malik, H. S. & Yao, M. C. 2016. The piggyBac transposon-derived genes TPB1 and TPB6 mediate essential transposon-like excision during the developmental rearrangement of key genes in Tetrahymena thermophila. Genes & Development, 30, 2724–2736.

David L. Spector, R. D. G., Leslie A. Leinwand 1998. *Cells: A Laboratory Manual, Volume 1,* Chapter 18, Culture and Manipulation of Tetrahymena.

Du, F., Edwards, K., Shen, Z., Sun, B., De Lozanne, A., Briggs, S. & Firtel, R. A. 2008. Regulation of contractile vacuole formation and activity in Dictyostelium. Embo J, 27, 2064–76.

Elliott, A. M. & Bak, I. J. 1964. The Contractile Vacuole and Related Structures in Tetrahymena Pyriformis. J Protozool, 11, 250–61.

Essid, M., Gopaldass, N., Yoshida, K., Merrifield, C. & Soldati, T. 2012. Rab8a regulates the exocyst-mediated kiss-and-run discharge of the Dictyostelium contractile vacuole. Mol Biol Cell, 23, 1267–82.

Fratti, R. A., Jun, Y., Merz, A. J., Margolis, N. & Wickner, W. 2004. Interdependent assembly of specific regulatory lipids and membrane fusion proteins into the vertex ring domain of docked vacuoles. J Cell Biol, 167, 1087–98.

Gabriel, D., Hacker, U., Kohler, J., Muller-Taubenberger, A., Schwartz, J. M., Westphal, M. & Gerisch, G. 1999. The contractile vacuole network of Dictyostelium as a distinct organelle: its dynamics visualized by a Gfp marker protein (vol 112, pg 3995, 1999). Journal of Cell Science, 112, U3–U3.

Gerald, N. J., Siano, M. & De Lozanne, A. 2002. The Dictyostelium Lvsa protein is localized on the contractile vacuole and is required for osmoregulation. Traffic, 3, 50–60.

Gorovsky, M. A., Yao, M. C., Keevert, J. B. & Pleger, G. L. 1975. Isolation of micro- and macronuclei of Tetrahymena pyriformis. Methods Cell Biol, 9, 311–27.

Harris, E., Yoshida, K., Cardelli, J. & Bush, J. 2001. Rab11-like GTPase associates with and regulates the structure and function of the contractile vacuole system in dictyostelium. J Cell Sci, 114, 3035–45.

Howard-Till, R. A. & Yao, M. C. 2006. Induction of gene silencing by hairpin Rna expression in Tetrahymena thermophila reveals a second small Rna pathway. Mol Cell Biol, 26, 8731–42.

Jimenez, V., Miranda, K. & Augusto, I. 2022. The old and the new about the contractile vacuole of Trypanosoma cruzi. J Eukaryot Microbiol, 69, e12939.

Kissmehl, R., Froissard, M., Plattner, H., Momayezi, M. & Cohen, J. 2002. Nsf regulates membrane traffic along multiple pathways in Paramecium. J Cell Sci, 115, 3935–46.

Klauke, N. & Plattner, H. 2000. “Frustrated Exocytosis”--a novel phenomenon: membrane fusion without contents release, followed by detachment and reajachment of dense core vesicles in Paramecium cells. J Membr Biol, 176, 237–48.

Klinger, C. M., Klute, M. J. & Dacks, J. B. 2013. Comparative genomic analysis of mul5-subunit tethering complexes demonstrates an ancient pan-eukaryotic complement and sculpting in Apicomplexa. PLos One, 8, e76278.

Ladenburger, E. M., Korn, I., Kasielke, N., Wassmer, T. & Plattner, H. 2006. An Ins(1,4,5)P3 receptor in Paramecium is associated with the osmoregulatory system. J Cell Sci, 119, 3705–17.

Ladenburger, E. M., Sehring, I. M., Korn, I. & Plattner, H. 2009. Novel types of Ca2+ release channels participate in the secretory cycle of Paramecium cells. Mol Cell Biol, 29, 3605–22.

Linkner, J., Witte, G., Zhao, H., Junemann, A., Nordholz, B., Runge-Wollmann, P., Lappalainen, P. & Faix, J. 2014. The inverse Bar domain protein IBARa drives membrane remodeling to control osmoregulation, phagocytosis and cytokinesis. J Cell Sci, 127, 1279–92.

Lorincz, P., Lakatos, Z., Varga, A., Maruzs, T., Simon-Vecsei, Z., Darula, Z., Benko, P., Csordas, G., Lippai, M., Ando, I., Hegedus, K., Medzihradszky, K. F., Takats, S. & Juhasz, G. 2016. Minicorvet is a Vps8-containing early endosomal tether in Drosophila. Elife, 5.

Macro, L., Jaiswal, J. K. & Simon, S. M. 2012. Dynamics of clathrin-mediated endocytosis and its requirement for organelle biogenesis in Dictyostelium. J Cell Sci, 125, 5721–32.

Malchow, D., Lusche, D. F., De Lozanne, A. & Schlatterer, C. 2008. A fast Ca2+-induced Ca2+-release mechanism in Dictyostelium discoideum. Cell Calcium, 43, 521–30.

Manna, P. T., Barlow, L. D., Ramirez-Macias, I., Herman, E. K. & Dacks, J. B. 2023. Endosomal vesicle fusion machinery is involved with the contractile vacuole in Dictyostelium discoideum. J Cell Sci, 136.

Morlon-Guyot, J., El Hajj, H., Martin, K., Fois, A., Carrillo, A., Berry, L., Burchmore, R., Meissner, M., Lebrun, M. & Daher, W. 2018. A proteomic analysis unravels novel Corvet and Hops proteins involved in Toxoplasma gondii secretory organelles biogenesis. Cell Microbiol, 20, e12870.

Naitoh, Y., Tominaga, T. & Allen, R. 1997. The contractile vacuole fluid discharge rate is determined by the vacuole size immediately before the start of discharge in Paramecium multimicronucleatum. J Exp Biol, 200, 1737–44.

Parkinson, K., Baines, A. E., Keller, T., Gruenheit, N., Bragg, L., North, R. A. & Thompson, C. R. 2014. Calcium-dependent regulation of Rab activation and vesicle fusion by an intracellular P2x ion channel. Nat Cell Biol, 16, 87–98.

Patel, S. & Docampo, R. 2010. Acidic calcium stores open for business: expanding the potential for intracellular Ca2+ signaling. Trends Cell Biol, 20, 277–86.

Peplowska, K., Markgraf, D. F., Ostrowicz, C. W., Bange, G. & Ungermann, C. 2007. The Corvet tethering complex interacts with the yeast Rab5 homolog Vps21 and is involved in endo-lysosomal biogenesis. Dev Cell, 12, 739–50.

Plattner, H. 2013. Contractile vacuole complex--its expanding protein inventory. Int Rev Cell Mol Biol, 306, 371–416.

Reuter, A. T., Stuermer, C. A. & Plattner, H. 2013. Identification, localization, and functional implications of the microdomain-forming stomatin family in the ciliated protozoan Paramecium tetraurelia. Eukaryot Cell, 12, 529–44.

Schindelin, J., Arganda-Carreras, I., Frise, E., Kaynig, V., Longair, M., Pietzsch, T., Preibisch, S., Rueden, C., Saalfeld, S., Schmid, B., Tinevez, J. Y., White, D. J., Hartenstein, V., Eliceiri, K., Tomancak, P. & Cardona, A. 2012. Fiji: an open-source planorm for biological-image analysis. Nat Methods, 9, 676–82.

Schonemann, B., Bledowski, A., Sehring, I. M. & Plattner, H. 2013. A set of Snare proteins in the contractile vacuole complex of Paramecium regulates cellular calcium tolerance and also contributes to organelle biogenesis. Cell Calcium, 53, 204–16.

Shang, Y., Song, X., Bowen, J., Corstanje, R., Gao, Y., Gaertig, J. & Gorovsky, M. A. 2002. A robust inducible-repressible promoter greatly facilitates gene knockouts, conditional expression, and overexpression of homologous and heterologous genes in Tetrahymena thermophila. Proc Natl Acad Sci U S A, 99, 3734–9.

Sivaramakrishnan, V. & Fountain, S. J. 2012. A mechanism of intracellular P2x receptor activation. J Biol Chem, 287, 28315–26.

Spang, A. 2016. Membrane Tethering Complexes in the Endosomal System. Front Cell Dev Biol, 4, 35.

Sparvoli, D., Richardson, E., Osakada, H., Lan, X., Iwamoto, M., Bowman, G. R., Kontur, C., Bourland, W. A., Lynn, D. H., Pritchard, J. K., Haraguchi, T., Dacks, J. B. & Turkewitz, A. P. 2018. Remodeling the Specificity of an Endosomal Corvet Tether Underlies Formation of Regulated Secretory Vesicles in the Ciliate Tetrahymena thermophila. Curr Biol, 28, 697–710 e13.

Sparvoli, D., Zoltner, M., Cheng, C. Y., Field, M. C. & Turkewitz, A. P. 2020. Diversification of Corvet tethers facilitates transport complexity in Tetrahymena thermophila. J Cell Sci, 133.

Stavrou, I. & O’halloran, T. J. 2006. The monomeric clathrin assembly protein, AP180, regulates contractile vacuole size in Dictyostelium discoideum. Mol Biol Cell, 17, 5381–9.

Stevens, T. H. & Forgac, M. 1997. Structure, function and regulation of the vacuolar (H+)-ATPase. Annu Rev Cell Dev Biol, 13, 779–808.

Tani, T., Tominaga, T., Allen, R. D. & Naitoh, Y. 2002. Development of periodic tension in the contractile vacuole complex membrane of paramecium governs its membrane dynamics. Cell Biol Int, 26, 853–60.

Tominaga, T., Allen, R. D. & Naitoh, Y. 1998a. Cyclic changes in the tension of the contractile vacuole complex membrane control its exocytotic cycle. J Exp Biol, 201 (Pt 18), 2647–58.

Tominaga, T., Allen, R. D. & Naitoh, Y. 1998b. Electrophysiology of the in situ contractile vacuole complex of Paramecium reveals its membrane dynamics and electrogenic site during osmoregulatory activity. J Exp Biol, 201, 451–60.

Ulrich, P. N., Jimenez, V., Park, M., Martins, V. P., Atwood, J., 3rd, Moles, K., Collins, D., Rohloff, P., Tarleton, R., Moreno, S. N., Orlando, R. & Docampo, R. 2011. Identification of contractile vacuole proteins in Trypanosoma cruzi. PLos One, 6, e18013.

Van Der Beek, J., Jonker, C., Van Der Welle, R., Liv, N. & Klumperman, J. 2019. Corvet, Chevi and Hops - multisubunit tethers of the endo-lysosomal system in health and disease. J Cell Sci, 132.

Wang, L., Merz, A. J., Collins, K. M. & Wickner, W. 2003. Hierarchy of protein assembly at the vertex ring domain for yeast vacuole docking and fusion. J Cell Biol, 160, 365–74.

Wen, Y., Stavrou, I., Bersuker, K., Brady, R. J., De Lozanne, A. & O’halloran, T. J. 2009. AP180-mediated trafficking of Vamp7b limits homotypic fusion of Dictyostelium contractile vacuoles. Mol Biol Cell, 20, 4278–88.

Yao, M. C. & Yao, C. H. 1991. Transformation of Tetrahymena to cycloheximide resistance with a ribosomal protein gene through sequence replacement. Proc Natl Acad Sci U S A, 88, 9493–7.

