## Supplementary figures and images for "VPS8D, a CORVET subunit, is required to maintain the contractile vacuole complex in *Tetrahymena thermophila*"

### Figure S1

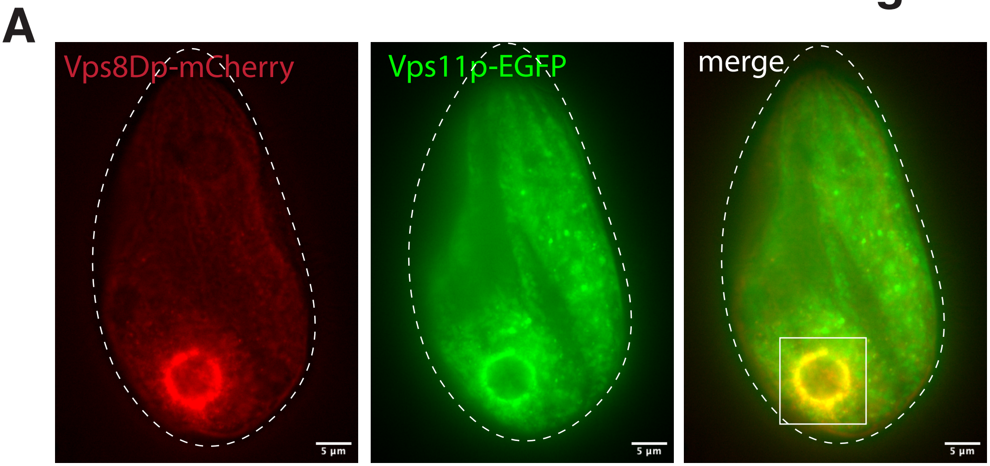

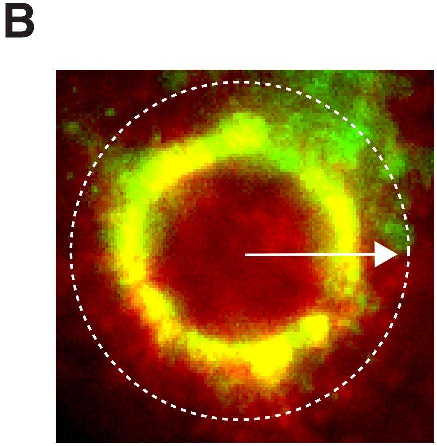

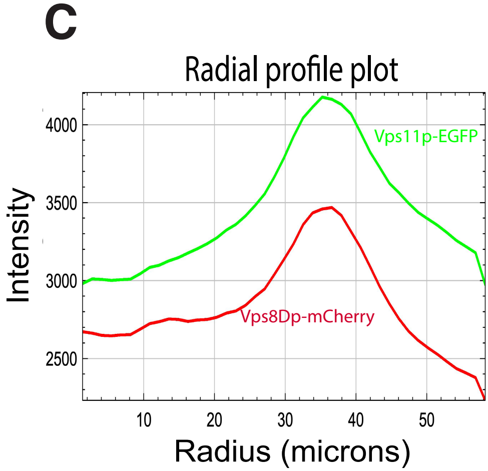

### Figure S2

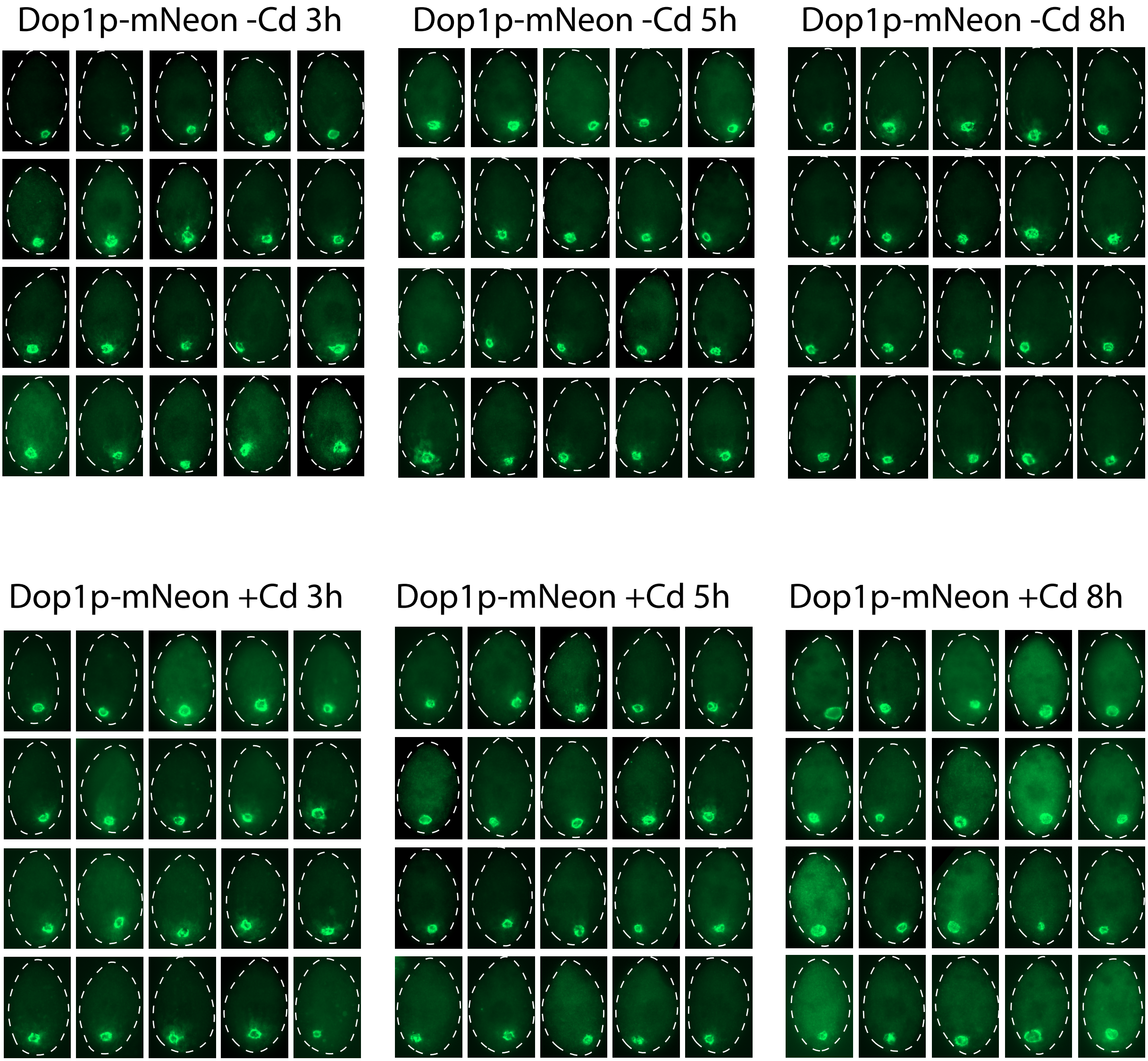

### Figure S3

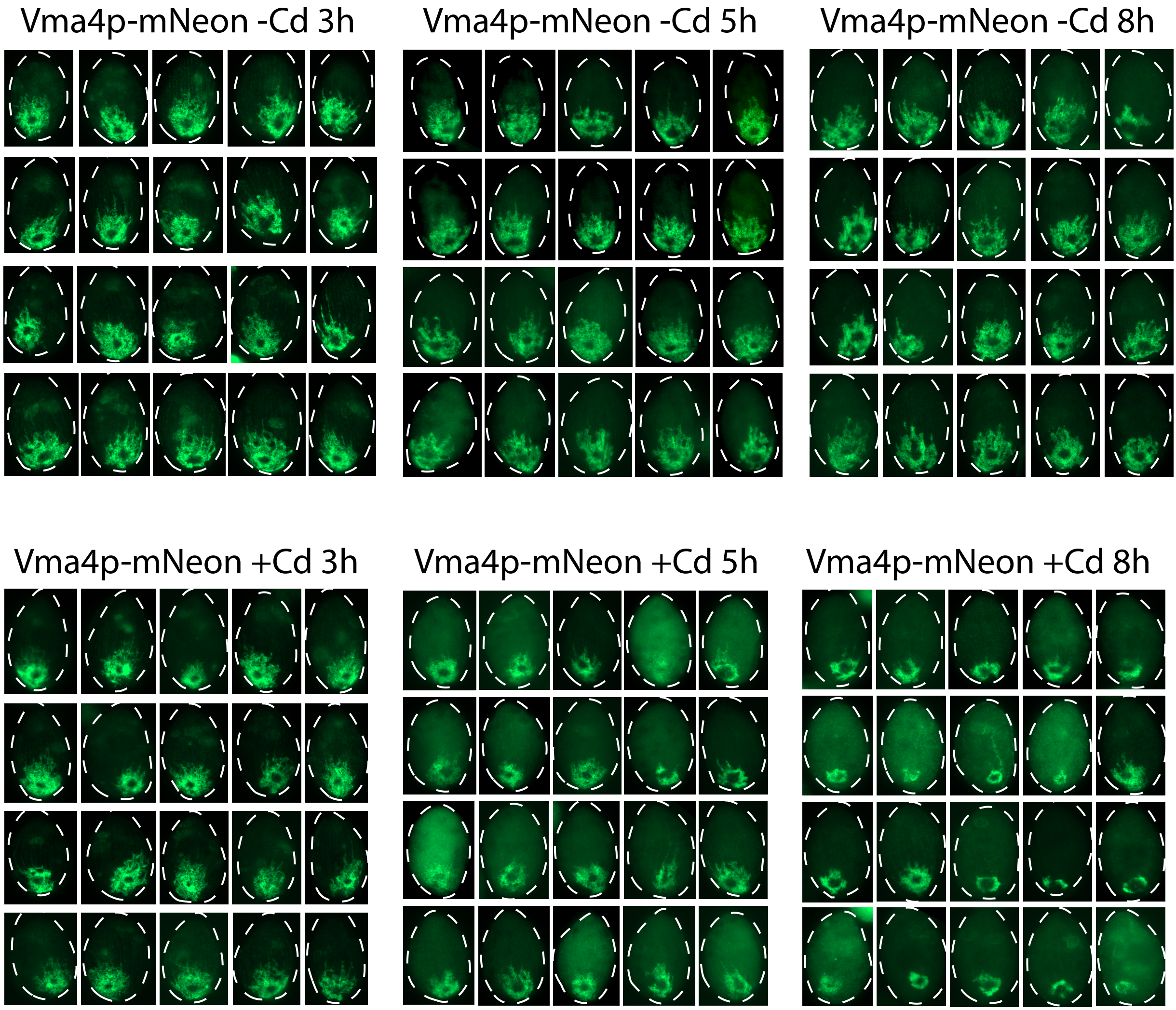

### Figure S4

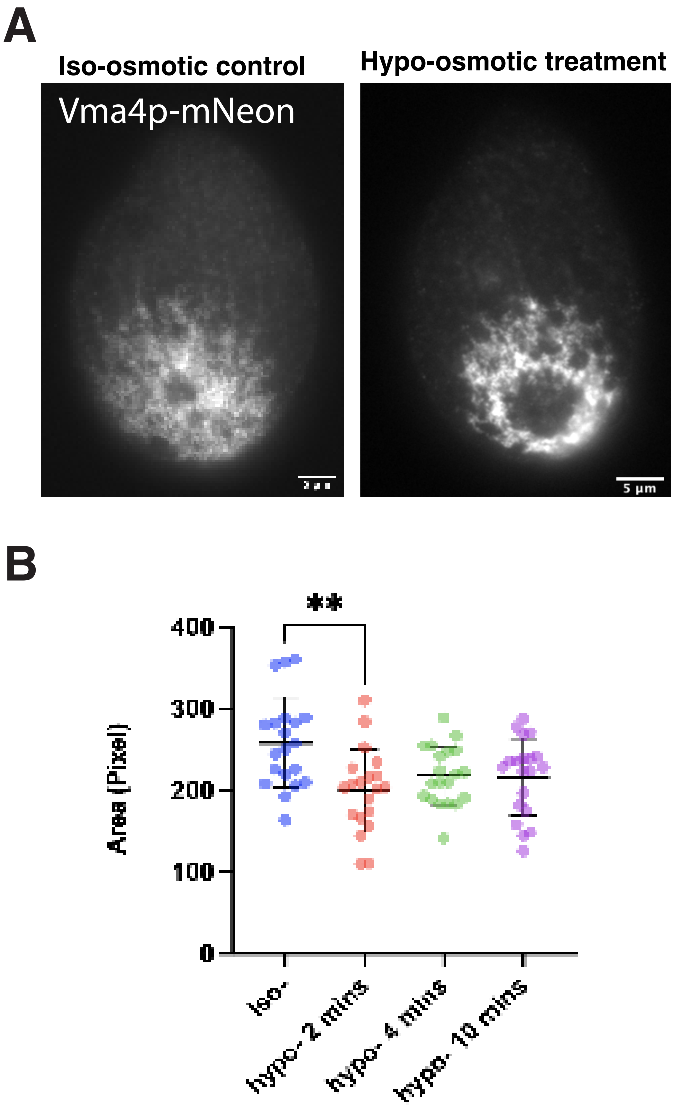

### Figure S5

#### Fig. S5

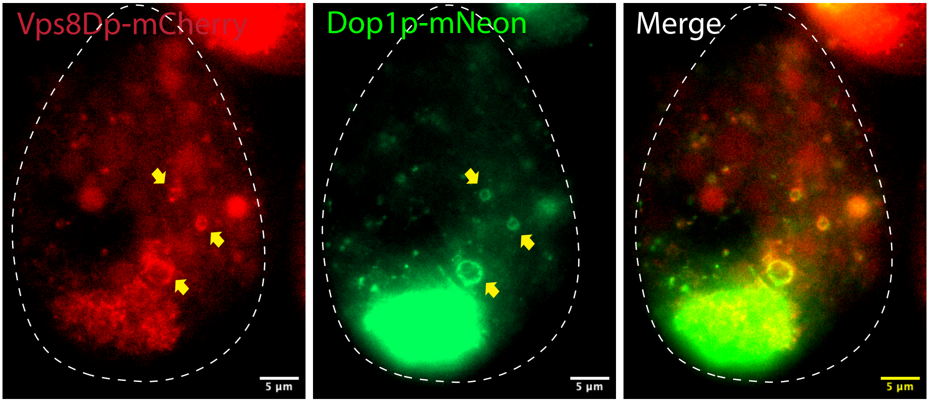
