## Supplementary material for "VPS8D, a CORVET subunit, is required to maintain the contractile vacuole complex in *Tetrahymena thermophila*": Supp Figure legend

Figure S1: Vps11p co-localizes with Vps8Dp at the bladder periphery.

A: Growing cell co-expressing Vps11-EGFP and Vps8Dp-mCherry was imaged for the mCherry channel in the left panel, GFP channel in the middle panel and the merge image in the right panel. Images were taken with a Marianas spinning disc confocal microscope.

B: The magnified image of boxed area from A containing the CV bladder. The fluorescent intensity profiles of the two fluorophores along the radial arrow shown were determined using the FIJI image Radial Profile plugin. See details in Materials and Methods of the previous study (reference).

C: The fluorescent intensity of Vps11p-EGFP and Vps8Dp-mCherry was plotted along the vector in B, drawn outward from the center of the CV bladder to the periphery. Vps11p and Vps8Dp show similar intensity profiles along a radial vector.

Figure S2: 20 images of VPS8Dhp -Cd and +Cd at 3h, 5h and 8h for the quantification of Dop1p at the CV bladder. VPS8Dhp cells expressing Dop1p-mNeon were grown for 3, 5 and 8 hours with and without adding cadmium, respectively and then were fixed for imaging. Images were taken with a Zeiss Axio Observer 7 system.

Figure S3: 20 images of VPS8Dhp -Cd and +Cd at 3h, 5h and 8h for the quantification of Vma4p-labelled spongiome. VPS8Dhp cells expressing Vma4p-mNeon were grown for 3, 5 and 8 hours with and without adding cadmium, respectively and then were fixed for imaging. Images were taken with a Zeiss Axio Observer 7 system.

Figure S4: The spongiome volume is reduced in hypoosmotic stressed cell.

A: Growing cells expressing Vma4p-mNeon in SPP medium for 16 hours as an iso-osmotic control sample (the image in the left panel). And cells were subjected to hypo-osmotic treatment for 2, 4 and 10 minutes as a hypo-osmotic stressed cell sample (the image in the right panel).

B: 20 images for each sample were analyzed to quantify the volume of Vma4p-labelled spongiome. The data were plotted using GraphPad Software Prism and using a two-tailed *t*-test. The mean±s.d. area (pixel) of Vma4p-labelled spongiome from each sample is shown. The difference between iso- and hypo- treatment for 2 mins condition is significant, *P*-value <0.05.

Figure S5: The Dop1p-labeled and Vps8Dp-labeled vacuoles appear to be the same structures in hypoosmotic stressed cell. The hypoosmotic stressed cell co-expressing Dop1p-mNeon and Vps8Dp-mCherry was imaged for the mCherry channel in the left panel, mNeon channel in the middle panel and the merge image in the right panel. Arrows point the large vesicles which are labeled by both Dop1p-mNeon and Vps8Dp-mCherry. Images were taken with a Zeiss Axio Observer 7 system.
