## Supplementary material for "VPS8D, a CORVET subunit, is required to maintain the contractile vacuole complex in *Tetrahymena thermophila*": Movie legend

Movie 1: FRAP analysis of wildtype cell expressing Dop1p-mNeon. Linked with Figure 2C. Live imaging with a Marianas spinning disc confocal microscope.

Movie 2: FRAP analysis of *VPS8D* knockdown cell expressing Dop1p-mNeon. Linked with Figure 2E. Live imaging with a Marianas spinning disc confocal microscope.

Movie 3: Live wildtype cell expressing Dop1p-mNeon with cross-sectional view of the CVC. Linked with Figure 4A. Live imaging with a Marianas spinning disc confocal microscope.

Movie 4: Live *VPS8D* knockdown cell expressing Dop1p-mNeon with cross-sectional view of the CVC. Linked with Figure 4B. Live imaging with a Marianas spinning disc confocal microscope.

Movie 5: Live *VPS8D* knockdown cell expressing Dop1p-mNeon with cross-sectional view of the CVC. Linked with Figure 4C. Live imaging with a Marianas spinning disc confocal microscope.

Movie 6: Live hypoosmotically stressed cell expressing Dop1p-mNeon with cross-sectional view of the CVC. Linked with Figure 5A. Live imaging with a Zeiss Axio Observer 7 system.

Movie 7: Live hyperosmotically stressed cell expressing Dop1p-mNeon with cross-sectional view of the CVC. Linked with Figure 5B. Live imaging with a Marianas spinning disc confocal microscope.

Movie 8: Live hypoosmotically stressed cell expressing Vps8Dp-mNeon with cross-sectional view of the CVC. Linked with Figure 5C. Live imaging with a Marianas spinning disc confocal microscope.

Movie 9: Large Vps8Dp-labeled vesicles that have leaked from ruptured hypoosmotically stressed cell expressing Vps8Dp-mNeon. Linked with Figure 5D. Live imaging with a Zeiss Axio Observer 7 system.

Movie 10: Large Vps8Dp-labeled vesicles that have leaked from a ruptured hypoosmotically stressed cell expressing Vps8Dp-mNeon. Vps8Dp redistributes to contact sites when two large vesicles come into contact. Linked with Figure 5E. Live imaging with a Zeiss Axio Observer 7 system.

Movie 11: Similar to Movie 10.
